# ISP Platform powered by Geneformer: Framework for Cross-Species, State-Feedback, and Multi-Gene *In Silico* Perturbation Screens with Application to iPS Cell State Transitions

**DOI:** 10.64898/2026.09.17.752497

**Authors:** Yuya Sanaki, Shuji Shigenobu, Ryusuke Niwa

## Abstract

*In silico* perturbation (ISP) enables virtual genetic screens by simulating targeted genetic manipulations in cells, prioritizing candidate genes before experiments. However, executing ISP requires programming expertise, limiting its use by experimental biologists. Here, we present the ISP Platform (https://github.com/YuyaSanaki/ISP-Platform, version 1.0.2), which automates ISP powered by Geneformer without code execution. The platform enables cross-species analyses in both directions by independently selecting input species and human or mouse Geneformer models. It also supports multi-gene ISP and implements State-feedback ISP, which predicts the gene expression ranking after each perturbation and feeds it into the next one. In single-cell transcriptomes of human and mouse induced pluripotent stem cells (iPSCs), genome-wide knockdown screens prioritized core pluripotency regulators with both species-matched and swapped models. *In silico* overexpression of seven pluripotency genes shifted somatic cells toward iPSCs more than the Yamanaka factors did. State-feedback ISP simulated all 24 orders of the Yamanaka factors and separated a factor-specific effect in every order, largest when POU5F1 or SOX2 came first. A pairwise *in silico* overexpression screen nominated candidate modulators of the primed-to-naïve transition beyond the NANOG and KLF2 benchmark. The ISP Platform makes Cross-species, State-feedback, and Multi-gene ISP available through a graphical user interface to guide experimental validation.

## 1. Introduction

Single-cell foundation models provide transferable representations for cell-state annotation, context-specific gene-network inference, and *in silico* perturbation (ISP; Ito et al., 2025; Kalfon et al., 2025; Theodoris et al., 2023). ISP computationally simulates genetic perturbations and predicts their effects on cellular state by quantifying cell-state shifts in the embedding space (Ito et al., 2025; Theodoris et al., 2023). Consequently, ISP offers an efficient framework to prioritize candidate regulators before conducting resource-intensive *in vivo* and *in vitro* screens. Following the development of the human Geneformer model (Theodoris et al., 2023) and, more recently, of Mouse-Geneformer (Ito et al., 2025), the utility of pretrained models has expanded across diverse biological contexts (Ito et al., 2025; Kobayashi et al., 2026; Theodoris et al., 2023). However, significant computational barriers (e.g., programming expertise, execution environments, and configuration) make it difficult for experimental biologists to access ISP without dedicated bioinformatics support.

Beyond simplifying computational accessibility, exploring how ISP can be aligned with complex biological contexts may provide a promising perspective. For instance, Geneformer systems currently exist as separate human and mouse models with distinct architectures (sequence length limits of 4,096 and 2,048 tokens; Chen et al., 2026; Ito et al., 2025). Ito et al. (2025) converted human genes to their mouse orthologs and applied Mouse-Geneformer to human scRNA-seq data. However, no framework for ISP has made the human and mouse models interchangeable in both directions, so that cells of either species can be analyzed with either model. Cross-species evaluation is practically valuable in translational workflows. Using mouse experimental data, it can assess whether candidate interventions induce the intended cell-state transitions in the human model. This allows researchers to evaluate functional conservation at the level of cellular phenotypes, rather than relying solely on gene orthology, thereby prioritizing candidates likely to be effective across species and reducing translational failures. Furthermore, complex cellular processes are rarely driven by isolated, single-gene events. Instead, these processes rely on multi-factorial regulatory programs acting in a temporally coordinated manner (Buganim et al., 2012; Polo et al., 2012).

To extend ISP, we developed the ISP Platform, an open-source framework powered by Geneformer designed to make advanced ISP accessible. The platform provides both a command-line and an interactive graphical user interface (GUI), enabling experimental biologists to perform reproducible ISP workflows without programming setup. To our knowledge, the ISP Platform is the first framework in which the human and mouse Geneformer models are interchangeable in both directions for ISP. To bridge the two models reliably, the platform integrates automated bidirectional orthology mapping that reports every gene dropped during conversion, an optional ortholog-loss quality control gate, and sequence-length normalization, which accommodates the architectural differences between the models. Furthermore, its perturbation engine applies length-preserving rank edits for multi-gene perturbations and feeds the model’s inferred response back between the steps of multi-step perturbations. All three types of ISP (Cross-species, State-feedback, and Multi-gene) can be run from the GUI.

Pluripotent stem cell biology provides an ideal testbed to benchmark these Cross-species, State-feedback, and Multi-gene perturbation capabilities. Human and mouse pluripotent stem cells share core regulatory circuitry but exhibit distinct transcriptional networks and stability conditions between primed and naïve states (Boyer et al., 2005; Gafni et al., 2013; Rodda et al., 2005; Takahashi et al., 2007; Takahashi & Yamanaka, 2006). Furthermore, cellular reprogramming into pluripotency is an inherently dynamic, temporally ordered process driven by defined transcription factors (Takahashi et al., 2007; Takahashi & Yamanaka, 2006). Leveraging the ISP Platform, which enables the decoupled selection of input species and model variants alongside automated orthology-aware configuration, we systematically evaluated these stem-cell transitions. Through Cross-species, State-feedback, and Multi-gene ISP screens, we prioritize candidate regulators of pluripotency maintenance and the primed-to-naïve transition, separate the factor-specific effects of the Yamanaka factors and of their order from placebo responses, and show that cross-species predictions recover core pluripotency regulators. Together, our platform provides an accessible and scalable framework to prioritize genetic interventions across complex biological programs.

## 2. Results

### 2.1 Scheme of the ISP Platform for Cross-species, State-feedback, and Multi-gene *in silico* perturbations

We developed the ISP Platform, a Geneformer-powered framework for virtual genetic screening that supports Cross-species, State-feedback, and Multi-gene *in silico* perturbation (ISP) analyses across diverse biological contexts (Fig. 1A). For cross-species ISP, users can independently specify the input single-cell transcriptomic dataset and Geneformer model variant (Fig. 1B). When these species differ, the platform converts cells to the model’s genes through bidirectional one-to-one Ensembl orthologs (human↔mouse), reports ortholog loss for QC, re-tokenizes with the model’s dictionary and input size (2,048 or 4,096 tokens), and fine-tunes the selected Geneformer model, so that ISP and goal states are evaluated in that model’s gene space (see Experimental Procedures).

**Fig. 1.**
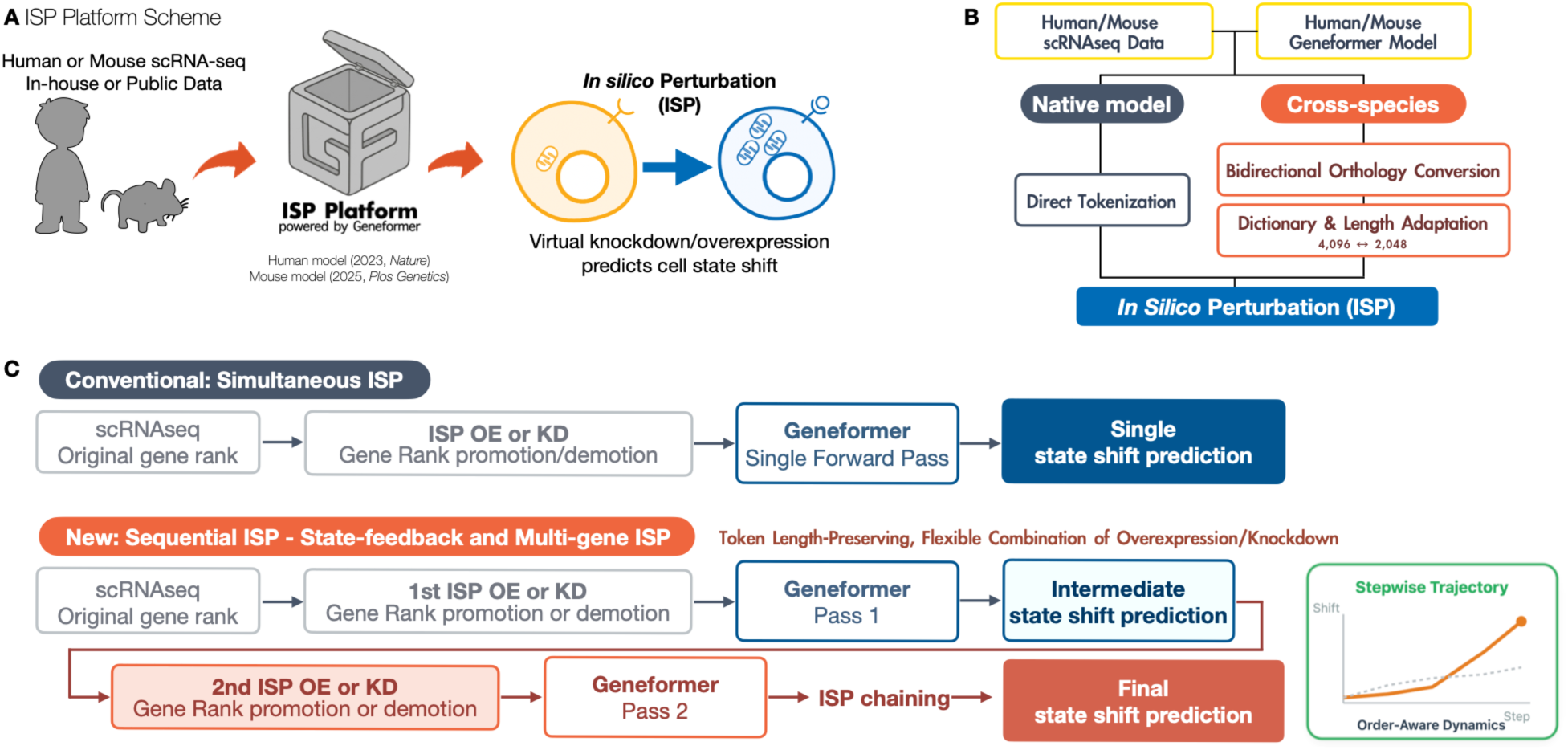
ISP Platform enables Cross-species, State-feedback, and Multi-gene ISP with an end-to-end pipeline. (A) Overview of the ISP Platform workflow. Human or mouse single-cell RNA-seq (scRNA-seq) data, in-house or public, are analysed with the human (Chen et al., 2026) or mouse (Ito et al., 2025) Geneformer model. Virtual knockdown (KD) or overexpression (OE) of genes predicts the shift of the cell state. (B) Data and model selection. The scRNA-seq data and the Geneformer model are chosen independently. With the model of the same species (native model), the cells are tokenized directly. With the model of the other species (cross-species ISP, swapped model), the genes are converted through one-to-one orthologs between human and mouse, and the cells are re-tokenized with the model’s dictionary and input size (4,096 tokens for the human model, 2,048 for the mouse model) before ISP. (C) State-feedback and multi-gene ISP. Length-preserving rank edits move overexpressed genes to the top rank and remove knocked-down genes. Without feedback, Geneformer only scores the edited encoding (top). In State-feedback ISP (bottom), the hidden-state change after each step is decoded into a reordering of the cell’s genes, which is the input to the next step. The goal-state shift is reported after every step.

To extend ISP beyond single-step perturbations, we implemented State-feedback ISP to simulate sequential perturbation (Fig. 1C). In State-feedback ISP, each perturbation step acts on the gene ranks that the model predicts after the preceding step. The simulation therefore takes the order of the perturbations into account, but not their timing. A step is not a unit of time, and the result is not a time course of the cell.

For multi-gene perturbations, the length of the encoding has to be kept constant. Overexpressing genes that are absent from a cell lengthens its rank-value encoding. Our length-preserving rank-editing strategy moves overexpressed genes that are already expressed to the front and, for each newly inserted gene, removes the lowest-ranked gene (Fig. 1C). Each cell therefore keeps its original token length, independently of the model’s input size. This approach enables flexible combinations of overexpression and knockdown perturbations.

### 2.2 Cross-species ISP screens prioritize conserved and species-specific regulators of pluripotency

To examine the platform capability, iPSC scRNA-seq datasets were used (Liu et al., 2020; Schiebinger et al., 2019). After predefined quality control and balanced subsampling (3,000 day-0 somatic and 3,000 iPS cells per arm, Supplementary Table 1), cells were tokenized with the selected dictionary and embedded using the corresponding fine-tuned checkpoint. Projecting human and mouse iPSCs alongside somatic counterparts demonstrated clear state separation under both matched and cross-species foundation models (Fig. 2A).

**Fig. 2.**
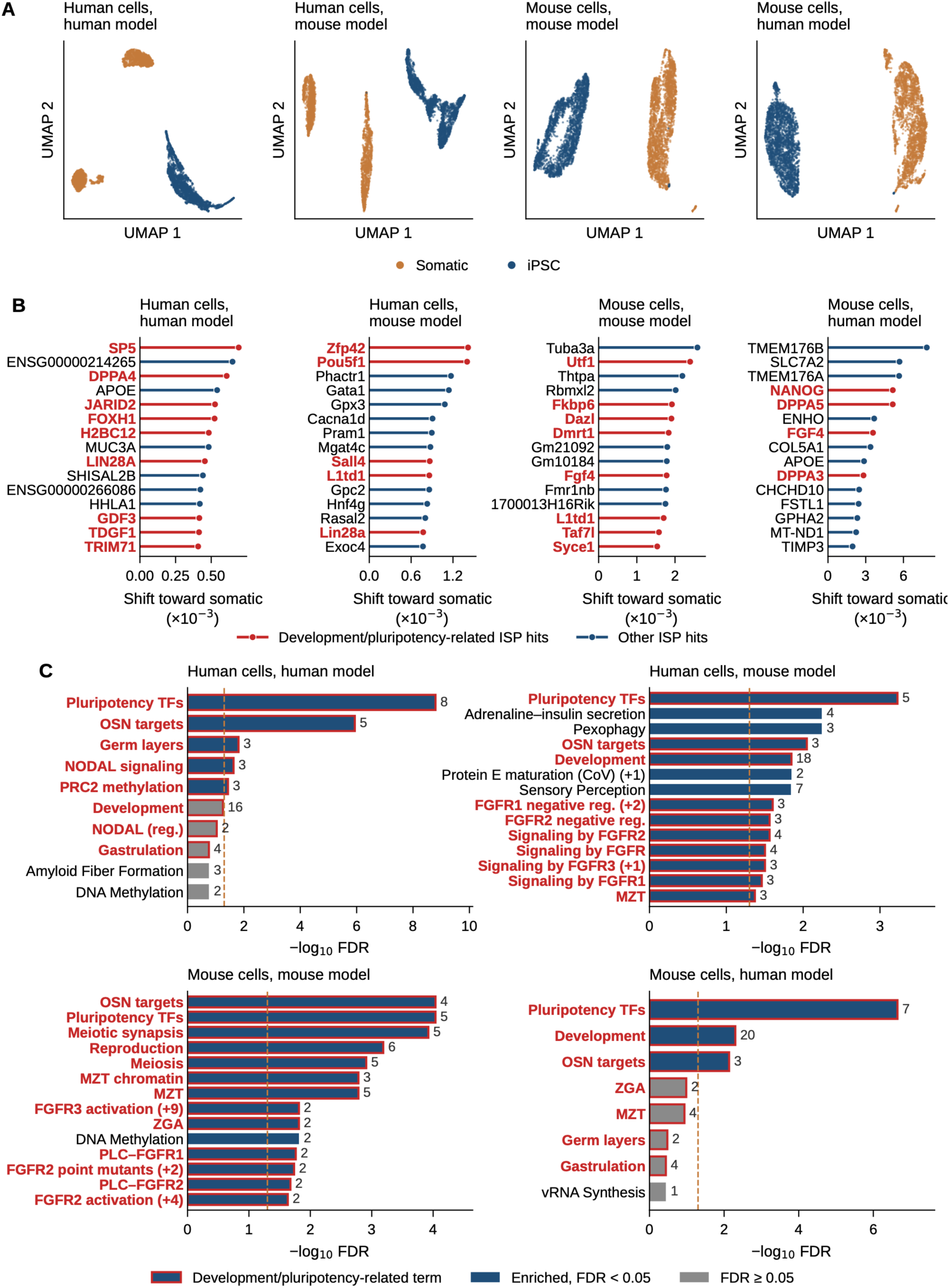
Cross-species genome-wide ISP screens prioritize pluripotency regulators across matched and swapped models. (A) UMAP projections of human and mouse iPSCs (blue) and their somatic counterparts (orange) embedded with the matched and the cross-species (swapped) fine-tuned models, with 3,000 cells per state in each arm. (B) Top 15 candidate regulators prioritized for pluripotency maintenance by genome-wide *in silico* knockdown (KD) screens from iPSCs toward the somatic state (FDR < 0.05, N_Detections > 20, the number of cells in which the gene was detected), ordered by the shift toward the somatic state. Known pluripotency- and development-associated genes are highlighted in red. (C) Reactome Pathways 2024 enrichment (Enrichr) of the top 100 genes of each screen (filters as in B). Shown are the 10 terms with the lowest FDR and further development- or pluripotency-related terms with FDR < 0.05, up to 14 rows. Bars, −log₁₀ FDR. Navy, FDR < 0.05. Grey, FDR ≥ 0.05. Dashed line, FDR = 0.05. Numbers, query genes in the term. Development- and pluripotency-related terms are outlined and labelled in red. Terms that contain the same query genes and have the same FDR are shown as one row, named after one of them, and (+n) gives the number of further terms in that row (for example, FGFR3 activation (+9) in mouse cells with the mouse model stands for ten FGFR terms that each contain only FGF17 and FGF4). Terms with equal FDR are shown or omitted together.

From the genome-wide *in silico* knockdown (KD) screens from pluripotent cells toward a somatic goal state across four matched and swapped-model arms, the top 15 candidate regulators are presented in Fig. 2B. Candidates were required to have FDR < 0.05 and to be detected in more than 20 of the 3,000 start cells (N_Detections > 20). N_Detections is the number of cells whose encoding contains the gene, and hence the number of per-cell shifts averaged for that gene. Because a gene can be knocked down only in cells that express it, this filter excludes genes whose shift rests on only a few cells. In the human-matched model, leading hits included established pluripotency and developmental regulators such as SP5, DPPA4, JARID2, and LIN28A. Similarly, the mouse-matched arm successfully retrieved known pluripotency-associated genes including Utf1 and Fgf4. Under model-swapping conditions, the prioritized sets retained several classical regulators, such as Zfp42, Pou5f1, Sall4, and Lin28a in the human-swapped arm and NANOG, DPPA5, FGF4, and DPPA3 in the mouse-swapped arm, alongside more heterogeneous hits (Fig. 2B).

Pathway enrichment analysis across the top 100 predicted genes further supported these findings (Fig. 2C). Reactome Pathways 2024 (Milacic et al., 2024) enrichment with Enrichr (Kuleshov et al., 2016; Xie et al., 2021) on each arm’s top 100 pluripotent→somatic genome-wide ISP hits (FDR < 0.05, N_Detections > 20) recovered coherent pluripotency programs in the human-matched arm, with Transcriptional regulation of pluripotent stem cells as the leading term and query genes including LIN28A, DPPA4, SALL4, ZSCAN10, PRDM14, NANOG, and TDGF1. Mouse-matched rankings showed robust enrichment for pluripotency transcriptional networks (DPPA4, SALL4, NANOG, TDGF1, and POU5F1), together with reproduction- and meiosis-related pathways. Transcriptional regulation of pluripotent stem cells was also the leading term in both swapped arms, with LIN28A, DPPA4, SALL4, ZSCAN10, PRDM14, NANOG, and POU5F1 in the mouse-swapped arm (mouse cells evaluated using the human model) and LIN28A, DPPA4, SALL4, NANOG, and POU5F1 in the human-swapped arm (human cells evaluated using the mouse model). This term was more significant with the human model than with the mouse model, both in human cells (FDR 1.6 × 10⁻⁹ and 5.9 × 10⁻⁴, respectively) and in mouse cells (FDR 2.3 × 10⁻⁷ and 9.2 × 10⁻⁵, respectively). In human cells evaluated with the mouse model, the next two terms (Pexophagy, and Adrenaline, noradrenaline inhibits insulin secretion) were unrelated to pluripotency (Fig. 2C).

Together, these results show that both species-matched and swapped models recover core pluripotency regulators. The human model prioritized pluripotency programs in mouse cells even more clearly than the mouse model did, whereas the mouse model gave less specific candidates for human cells.

### 2.3 Multi-gene ISP of reprogramming factors

Somatic cell reprogramming is classically induced by overexpression (OE) of the Yamanaka factors: POU5F1 (OCT4), SOX2, KLF4, and MYC (OSKM; Takahashi et al., 2007; Takahashi & Yamanaka, 2006). In the original Geneformer study, simultaneous *in silico* OE of OSKM in fibroblasts shifted gene embeddings toward the iPSC state (Theodoris et al., 2023). In the ISP Platform, multi-gene OE uses a length-preserving rank edit (Fig. 3A). An OE gene already expressed in the cell is moved to the front, each absent OE gene is inserted at the front, and each inserted gene displaces the lowest-ranked gene, so that the perturbed encoding keeps the length of the cell. Both encodings are embedded with the OE genes included, and the shift is computed from the genes present in both, so the procedure is the same in every cell and at any model input size. We first applied simultaneous *in silico* OE of OSKM to human somatic cells using the human-matched model. Simultaneous OSKM OE produced a modest median goal-state shift toward the iPSC target (+0.0103, Fig. 3B). Because Yamanaka factors primarily initiate reprogramming through transcriptional and epigenetic remodeling to activate the endogenous pluripotency networks (Buganim et al., 2012; Polo et al., 2012), we reasoned that direct overexpression of endogenous core pluripotency factors POU5F1, SOX2, and NANOG (OSN; Boyer et al., 2005; Rodda et al., 2005) might drive a more pronounced shift. Indeed, simultaneous *in silico* OE of OSN produced a median shift of +0.0153 (n = 1,000, fraction of positive cells = 1.0), significantly exceeding that of OSKM under identical conditions (Fig. 3B). We then tested an expanded pluripotency-associated factor set comprising NANOG, POU5F1, SOX2, ESRRB, LIN28A, DPPA4, and TERT (the 7-factor cocktail; Boyer et al., 2005; Feng et al., 2009; Hernandez et al., 2018; Kinoshita et al., 2014; Yu et al., 2007). This 7-factor cocktail further elevated the median shift to +0.033 (n = 1,000, fraction of positive cells = 1.0), significantly outperforming both OSKM and OSN (Fig. 3B). These results suggest that Yamanaka factors primarily trigger reprogramming rather than directly establish the pluripotent state, suggesting that ISP evaluates cell-state shifts relative to target embeddings rather than predicting complex, downstream biological outcomes.

**Fig. 3.**
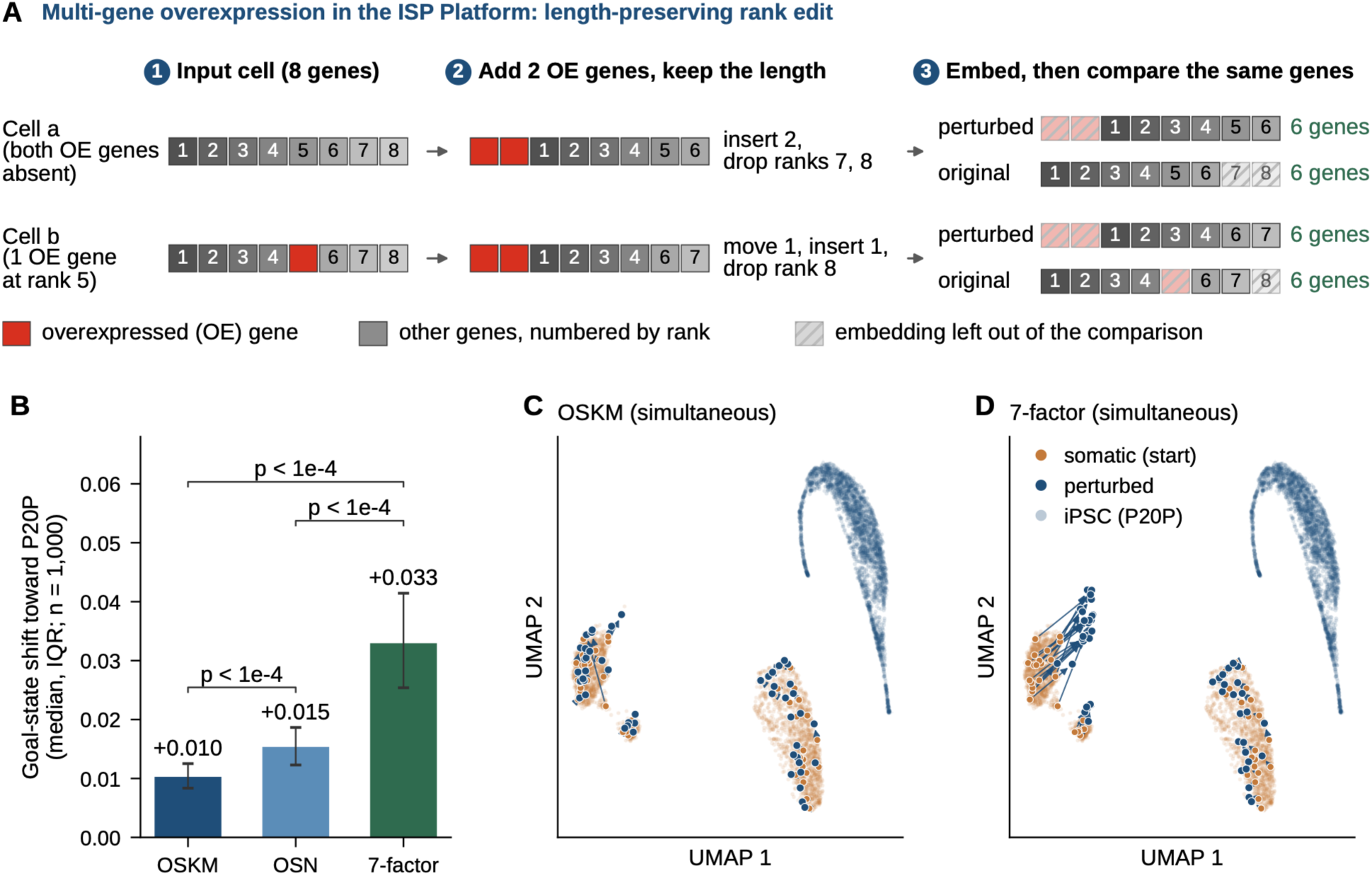
Multi-gene ISP of reprogramming factors. (A) Multi-gene overexpression (OE) in the ISP Platform (length-preserving rank edit), shown for two OE genes and two cells of eight genes. Boxes are the genes of a cell in rank-value order (numbers, expression rank in the original cell), and red boxes are OE genes. (1) In cell a both OE genes are absent, whereas in cell b one of them is already expressed (rank 5). (2) An OE gene already present is moved to the front, each absent one is inserted at the front, and each inserted gene displaces the lowest-ranked gene, so the perturbed encoding keeps the cell’s length. (3) Geneformer embeds the original and the perturbed encoding as they are, with the OE genes included, so the OE genes change the embeddings of all other genes through attention. The OE genes have no counterpart in the original cell. Their embeddings, and in the original those of the OE gene already present (cell b) and of the displaced genes, are left out (hatched). The embeddings of the same six genes are then compared gene by gene, or mean-pooled for the shift of the cell toward the goal state. The procedure is the same in every cell and at any model input size. (B) Goal-state shift of somatic cells toward P20P iPSCs after simultaneous overexpression of POU5F1, SOX2, KLF4 and MYC (OSKM), of POU5F1, SOX2 and NANOG (OSN) and of the 7-factor cocktail (NANOG, POU5F1, SOX2, ESRRB, LIN28A, DPPA4 and TERT) (median and IQR, n = 1,000, two-sided paired Wilcoxon test with Bonferroni correction). (C, D) Joint UMAP of somatic cells (orange, start), P20P iPSCs (light grey) and the same somatic cells after simultaneous overexpression (navy) of OSKM (C) or of the 7-factor cocktail (D). Lines show the displacement of 50 randomly chosen cells. Each group has 3,000 cells, which differ from those in (B).

In the embedding space, OSKM overexpression displaced somatic cells only slightly, whereas overexpression of the 7-factor cocktail displaced them toward the iPSC embeddings (Fig. 3C, D).

### 2.4 State-feedback ISP separates factor-specific effects of multi-step perturbations from placebo responses

Cellular reprogramming proceeds through sequential changes in gene expression, in which earlier changes alter the cell state on which later ones act (Buganim et al., 2012; Polo et al., 2012). Single-step ISP edits the encoding once and scores the result, so it cannot represent such sequential changes. To mimic sequential cell-state changes, we implemented State-feedback ISP, in which the predicted response to one perturbation becomes the input to the next (Fig. 1C). Geneformer, however, predicts changes in its hidden states and has no output for gene expression ranking. We therefore built a decoder that converts the model’s response into a new gene order. For each gene, a single linear layer translates the change in its final-layer hidden state into a shift of its rank, and the genes of the cell are reordered accordingly. The decoder is trained on the rank displacement of each gene between somatic cells and iPSCs (see Experimental Procedures), and the reordered encoding is the input to the next perturbation step.

We applied State-feedback ISP to the Yamanaka factors in one fixed order, adding one factor per step (KLF4→MYC→SOX2→POU5F1, K→M→S→O) to 300 somatic cells. All 24 orders are compared below. To test whether the decoded shift depended on the factors, we compared OSKM with placebo gene sets. A placebo gene set consists of genes drawn at random from the model’s vocabulary, each matched to one factor for detection rate and expression rank in the start cells (Supplementary Table 2). Genes specified for OE or KD are never used as placebo genes. A placebo gene therefore receives the same kind of rank edit as the factor it replaces, but it is chosen without regard to its biological function. We measured the effect of each gene set as its feedback gain, the additional shift toward the goal state compared with the same steps applied without feedback. Without correction, the placebo gene sets reached 95–96% of the feedback gain of OSKM (Fig. 4A). Most of the decoded shift therefore did not depend on which genes were added.This non-specific shift arises because rank editing alters Geneformer’s hidden states in a similar way regardless of the edited genes, and the decoder, trained on the transition from the start to the goal state, translates this change into a shift toward the goal state. Because the decoded shift is fed back after every step, the non-specific shift accumulates.

**Fig. 4.**
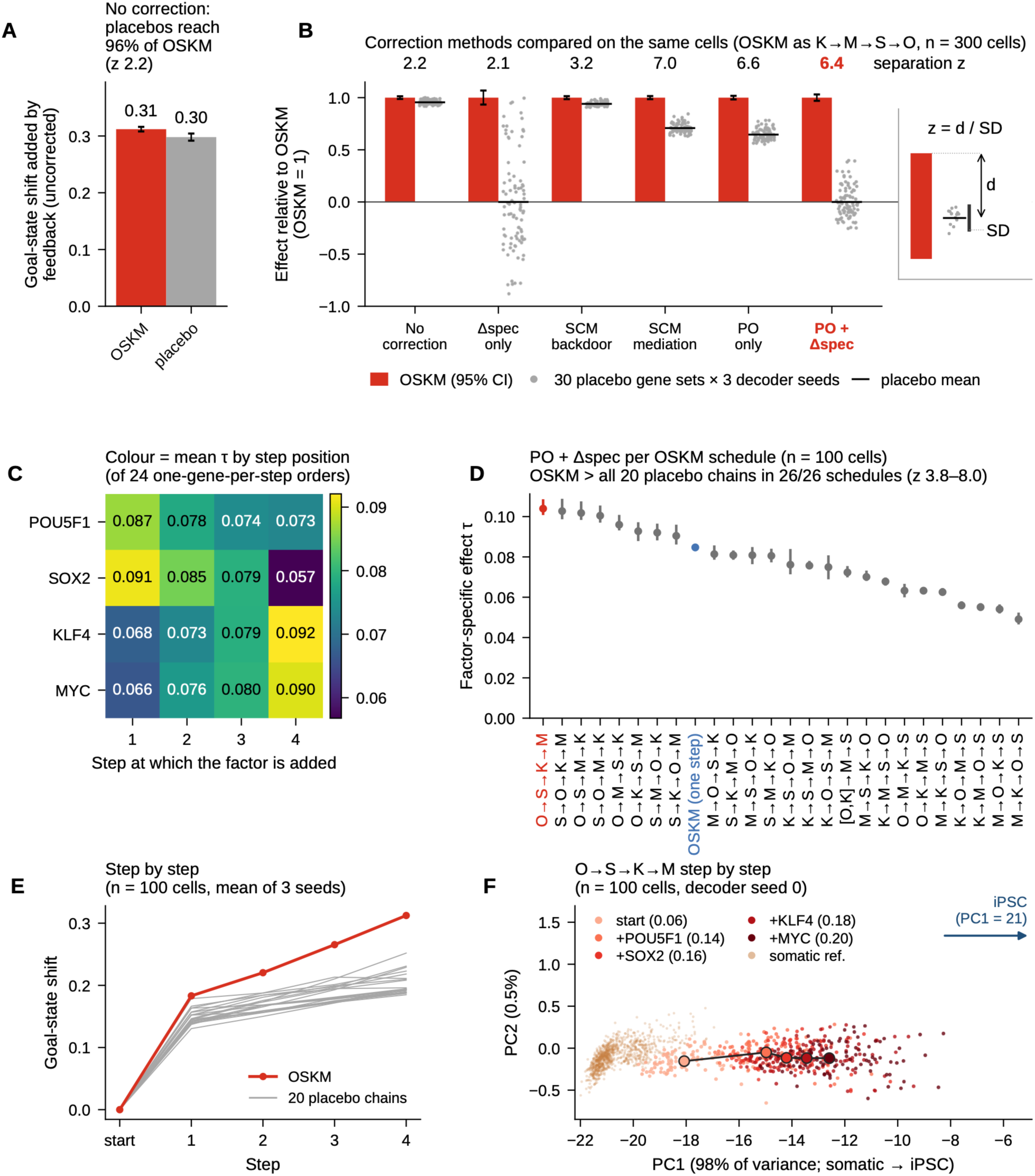
State-feedback ISP: separation from placebo responses and order of the Yamanaka factors. (A, B) OSKM added one factor per step in the order KLF4→MYC→SOX2→POU5F1 (K→M→S→O) in 300 somatic cells, compared with 30 placebo sets of four genes. Each placebo gene is matched to one factor for detection and rank in the start cells and added at that factor’s step (Fig. 1C). The same sets are used for all three decoder seeds. Bars, mean over seeds. Error bars, 95% CI (cell bootstrap) for OSKM and mean ± SD for the placebos. (A) Without correction, the goal-state shift added by feedback (end point with minus without feedback) is 96% as large for the placebos as for OSKM. (B) Correction methods, with OSKM set to 1: none, Δspec (feedback gain of OSKM minus the mean placebo gain, τ), structural causal model with backdoor or mediator adjustment, potential outcome (PO, each step decodes the hidden-state change of OSKM minus that of the placebos in the same cell), and PO + Δspec, used in (C–F). Dots, the 30 placebo sets with each seed, relative to OSKM of the same seed. Lines, their mean (0 by definition with Δspec). Numbers, z = d / SD, with d the distance of OSKM from the placebo mean and SD the standard deviation of the placebos (mean over seeds). (C, D) τ with PO + Δspec in 100 somatic cells. Each schedule is compared with its own 20 placebo chains, in which each factor is replaced at its step by a matched placebo gene. τ is smaller than in (A, B) because fewer cells are used, so schedules are compared only within (C, D). (C) Mean τ of the six one-gene-per-step orders that add each factor (row) at each step (column). (D) τ of the 24 one-gene-per-step orders, [POU5F1, KLF4]→MYC→SOX2 (OK→M→S, Liu et al., 2013) and one-step OSKM (blue). Red, POU5F1→SOX2→KLF4→MYC, the top-ranked order, followed in (E, F). Dots, mean of three seeds. Lines, range over seeds. OSKM exceeded every placebo chain in all schedules and seeds. (E) Goal-state shift after each step of this order (red) and of its 20 placebo chains (grey), mean of three seeds. (F) The cells of (E) at the start and after each feedback step (seed 0, n = 100), projected onto the first two principal components of 1,000 reference somatic cells and 1,000 reference iPSCs (Geneformer embeddings). PC1 (98% of the variance) separates the two. The iPSCs lie beyond the view at PC1 ≈ 21 (arrow). Large dots and line, median of each step. Numbers in brackets, mean position from the somatic (0) to the iPSC centroid (1) in the full embedding.

We then compared corrections that remove the non-specific shift at different points, using *in silico* OSKM overexpression as a benchmark (see Experimental Procedures). Three corrections act on the hidden-state change at every feedback step, before decoding. Under a structural causal model (SCM), backdoor adjustment removes baseline bias from each gene’s start-encoding hidden state. A regularized linear model (ridge regression) is fitted on correction placebos to learn how the hidden-state change depends on that start state. Its prediction is then subtracted from the observed change before decoding. In contrast, SCM mediator adjustment removes the non-specific response to the rank edit itself, which arises regardless of which genes are edited. At each step, the factors are swapped for matched placebo genes in the current encoding, and the mean hidden-state change across these placebos estimates this generic artifact, which is subtracted from the factor-induced change before decoding with the uncorrected decoder. The potential-outcome (PO) contrast subtracts the same placebo change but uses a decoder trained on this contrast. The gene-specific gain (Δspec, reported as τ) instead acts on the outcome and subtracts the mean feedback gain of placebo gene sets from that of the factors. Separation was measured as z, the distance of OSKM from the mean of the placebo sets in units of their standard deviation (Fig. 4B). Backdoor adjustment yielded weaker separation (z = 3.2, Fig. 4B). In contrast, the PO contrast and the SCM mediator adjustment, which directly subtract the response to the placebo sets, separated the effect of OSKM clearly (z = 6.6 and 7.0, respectively, Fig. 4B).

Because a gene-independent placebo response partially persisted under both approaches (placebo gene sets reached 64–65% and 70–72% of the OSKM gain, respectively, Fig. 4B), we established a two-tier correction workflow based on the potential-outcome framework. State-feedback ISP allocates expression-matched placebo gene sets into two groups: an internal group of 10 correction placebo gene sets and an external group of 30 comparison placebo gene sets. Both groups are chosen automatically from the start cells of the input dataset. Within each perturbation step, the correction placebos are swapped in for the query genes in the current encoding, and their mean hidden-state shift is subtracted from that of the query genes before decoding and feedback. In parallel, the comparison placebos are run through the entire simulation pipeline under identical conditions to establish an empirical null distribution. After the final step, Δspec (τ) is calculated by subtracting the mean feedback gain of the comparison placebos from that of the query genes.

Because the comparison placebos are never used for the correction, the perturbation is not evaluated against the same sets that corrected it. With this two-tier correction, the specific effect of OSKM was τ = 0.127–0.132 across three decoder seeds (95% CI 0.123–0.136), and OSKM ranked first among the 30 comparison placebos in every seed (z = 6.3–6.5, Fig. 4B). Thus, the two-tier correction achieved separation comparable to the single corrections while removing the placebo response from the estimate (Fig. 4B).

We next asked whether the order in which the Yamanaka factors are added changes their factor-specific effect. In 100 somatic cells, we evaluated all 24 orders that add one factor per step, the schedule [POU5F1, KLF4]→MYC→SOX2, which outperformed simultaneous delivery in mouse fibroblasts (Liu et al., 2013), and simultaneous OSKM. Each schedule was compared with its own 20 placebo chains, in which every factor was replaced, in the same step, by a matched placebo gene. The effect depended on the order (τ = 0.049–0.104), and the ranking of the orders was consistent across decoder seeds (Spearman 0.98–0.99). The effect was largest when POU5F1 or SOX2 was added first and KLF4 or MYC last (Fig. 4C). OSKM exceeded every placebo chain in all schedules and decoder seeds (z = 3.8–8.0, Fig. 4D).

Simultaneous OSKM ranked 9th of the 26 schedules (τ = 0.085). The eight orders that began with POU5F1 or SOX2 and ended with KLF4 or MYC exceeded it, and the other 16 fell below it (paired Wilcoxon signed-rank test with Holm correction, Fig. 4D). The schedule [POU5F1, KLF4]→MYC→SOX2 ranked 17th and fell below simultaneous OSKM in 93% of cells.

Followed step by step, the highest-ranked order, POU5F1→SOX2→KLF4→MYC, moved the cells further toward the goal state than every placebo chain from the first step on, in every decoder seed (Fig. 4E). To visualize these steps, we projected the cells onto the embedding space of reference somatic cells and iPSCs. Each step moved the cells further toward the iPSCs: the cells started at 6% of the distance between the somatic and iPSC centroids and reached 14%, 16%, 18% and 20% after the four steps. Every cell moved closer to the iPSCs at every step (Fig. 4F).

### 2.5 Targeted NANOG-plus-partner overexpression predicts modulators of the primed-to-naïve transition

Human pluripotent stem cells can be maintained in naïve or primed states with distinct molecular and developmental properties, corresponding to pre- and post-implantation epiblast states (Gafni et al., 2013; Weinberger et al., 2016). By contrast, mouse embryonic stem cells maintained in 2i/LIF culture conditions adopt a ground state that functionally parallels human naïve rather than human primed pluripotency. Building on this species-specific framework, we applied targeted two-gene OE screens from primed human P20P cells toward either the P20N naïve goal state (P20P→P20N) or the mouse DiPSC_2i 2i-maintained ground-state reference embedded in the human model (P20P→DiPSC_2i). Because the combination of NANOG and KLF2 was previously shown to facilitate primed-to-naïve transition (Takashima et al., 2014), we used NANOG+KLF2 as a positive benchmark and the mean shift across NANOG paired with negative-control partners as the baseline aggregate (Fig. 5, Supplementary Table 1). All screens used the human model, which was fine-tuned separately for each route on its start and goal cells.

**Fig. 5.**
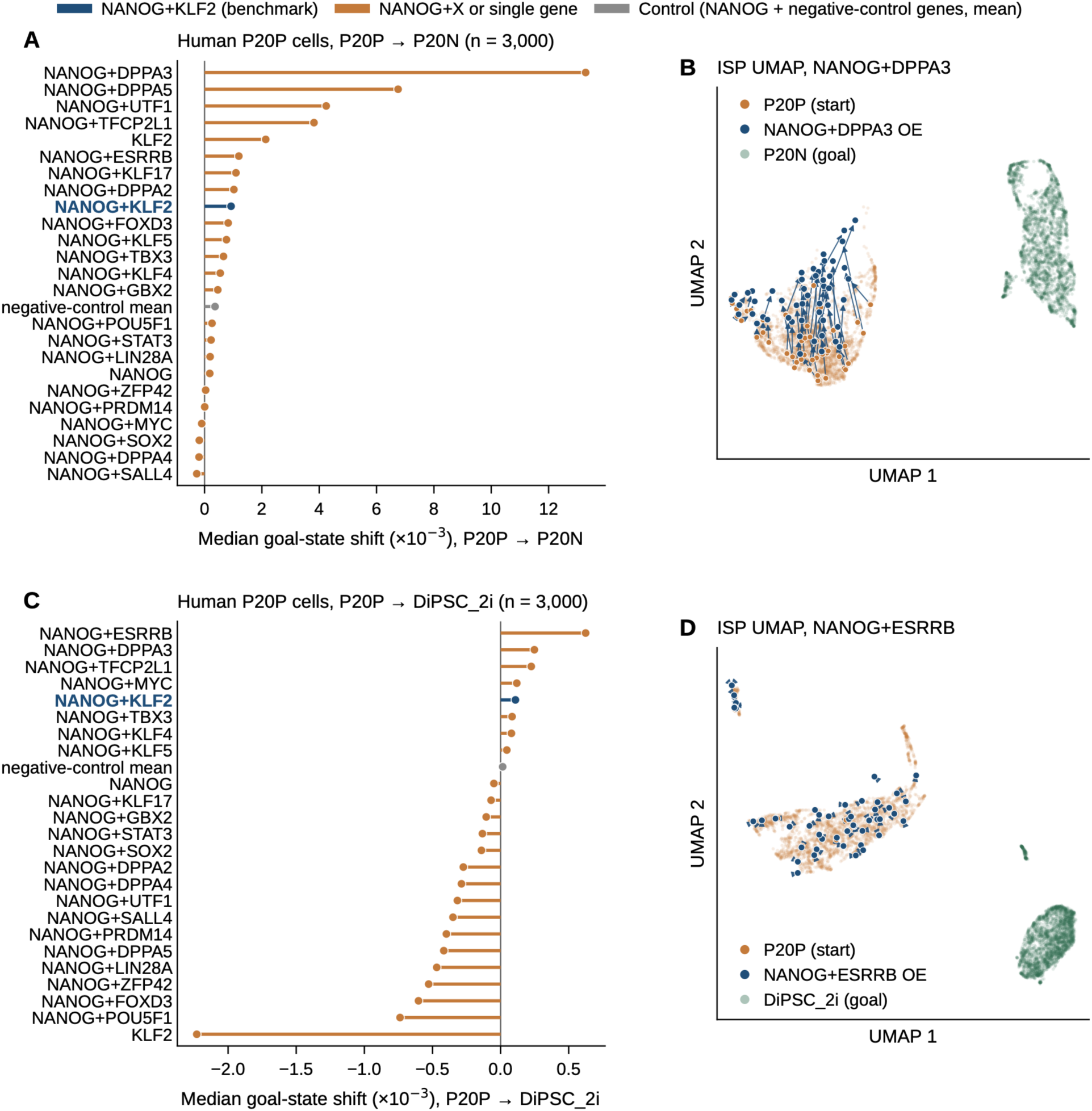
Cross-species ISP prioritizes pairwise regulators for primed-to-naïve pluripotency transitions. (A, B) Targeted two-gene overexpression screen from human primed iPSCs (P20P) toward naïve iPSCs (P20N). (A) Median goal-state shift (×10^−3^) after *in silico* overexpression of NANOG with each of 21 candidate partners and of NANOG or KLF2 alone (orange). Navy, the benchmark NANOG + KLF2 (Takashima et al., 2014). Grey, mean of the median shifts of NANOG paired with each of four negative-control genes (DDO, KRTAP12-3, ASIC5, PDHA2). Each perturbation was evaluated in 3,000 cells. (B) Joint ISP UMAP of P20P cells (orange, start), the same cells after overexpression of NANOG + DPPA3, the top partner in (A) (navy), and P20N cells (green, goal), with 3,000 cells per group. Lines show the displacement of 50 randomly chosen cells. (C, D) The same screen toward mouse ground-state iPSCs maintained in 2i (DiPSC_2i), embedded in the human model. (C) As in (A). (D) As in (B) for NANOG + ESRRB, the top partner in (C), with DiPSC_2i cells as the goal. Shifts are compared within a route. Their size differs between routes because the goal embeddings differ. The screens test selected partners, not the whole genome.

On the P20P→P20N route, seven partners exceeded the NANOG+KLF2 benchmark median shift (+0.00092), led by NANOG+DPPA3 (median = +0.0133), followed by DPPA5, UTF1, TFCP2L1, ESRRB, KLF17, and DPPA2 (Fig. 5A). NANOG alone and several core pluripotency pairings (for example NANOG+SOX2 or NANOG+SALL4) did not exceed the benchmark under this goal embedding, indicating that partner identity shapes the predicted shift. Several top partners are naïve-associated transcription factors or epigenetic regulators, supporting the prioritization of candidates beyond the NANOG/KLF2 benchmark for embedding shifts from the primed toward the naïve state. The ISP UMAP of the top P20N-route partner illustrated a clear displacement of P20P cells along the defined goal axis relative to unperturbed controls (Fig. 5B).

On the P20P→DiPSC_2i route, NANOG+ESRRB (median = +0.00062) exceeded the route-specific NANOG+KLF2 benchmark (median = +0.00011), as did NANOG+DPPA3, NANOG+TFCP2L1, and NANOG+MYC (Fig. 5C). For the *in silico* OE of NANOG+ESRRB, the joint ISP UMAP showed little displacement from the original primed state (Fig. 5D), consistent with the much smaller shifts than on the P20N route. Predicted partner rankings toward the human naïve state and the mouse ground-state reference were only moderately concordant (Spearman ρ ≈ 0.35 among shared partners). DPPA3 and TFCP2L1 ranked high on both routes, whereas DPPA5 and UTF1, second and third on the P20N route, fell below the negative-control mean toward DiPSC_2i, and ESRRB moved from fifth among the partners on the P20N route to first toward DiPSC_2i. Because the prioritized partner genes differed between the human naïve and mouse ground-state targets, these divergent predictions are consistent with differences between the defined human naïve and mouse 2i-reference embedding goals, while model- and mapping-related effects cannot be excluded.

## 3. Discussion

The ISP Platform provides a Docker-containerized framework that integrates tokenization, fine-tuning, and ISP while allowing human and mouse Geneformer models to be selected independently of input species. Whereas Mouse-Geneformer was applied to human cells after ortholog conversion (Ito et al., 2025), the platform makes the two models interchangeable in both directions. Cross-species analyses incorporate orthology-aware conversion, model-specific dictionaries and sequence-length constraints, and pre-tokenization reporting of mapping yield. The platform also supports simultaneous multi-gene ISP and multi-step ISP with feedback of the model’s response (State-feedback ISP), and all three types of ISP can be run from a graphical user interface.

Genome-wide knockdown prioritized genes whose removal shifted pluripotent cells toward somatic embeddings, and core pluripotency regulators were recovered with both species-matched and swapped models (Fig. 2). How far cross-species predictions can be relied on depended on the direction. In mouse cells, the pluripotency terms were more significant with the human model than with the mouse model itself (Fig. 2C), so mouse experimental data can be examined in the larger human model. In human cells evaluated with the mouse model, by contrast, the pluripotency terms were less significant and were interspersed with terms unrelated to pluripotency (Fig. 2C). Because the human model gave the more significant terms in the cells of both species, this asymmetry is more likely to reflect the models than the species of the cells. The human model was pretrained on more cells and has more parameters than the mouse model (Chen et al., 2026; Ito et al., 2025).

Converting human cells for the mouse model also retained fewer genes than the reverse conversion, and the mouse model accepts only half as many input genes. Cross-species predictions therefore depend on the model and its training data, so they prioritize candidates for experimental validation and do not replace it. We regard the species-matched model as the default and use cross-species ISP for three purposes: to borrow a model trained on a larger or more relevant corpus, to compare cells with a reference state available only in the other species, and to identify candidates that are robust to the choice of model. Candidates supported by both models and pathway-level agreement are therefore more informative than raw shift magnitudes, and we compare ranking overlap and pathway coherence within each input species rather than raw shift magnitudes across models. Genes without one-to-one orthologs can be evaluated in the swapped arms only through a curated ortholog table, as done here for POU5F1 and NANOG.

In multi-gene ISP, the 7-factor cocktail of pluripotency-associated genes shifted somatic cells further toward iPSCs than OSKM did (+0.033 vs +0.010, Fig. 3B). Ultimately, the ISP Platform provides an efficient, hypothesis-generating environment to benchmark multi-gene and multi-step perturbation programs, prioritizing candidate combinations for focused experimental validation.

Standard ISP evaluates single-gene or simultaneous multi-gene perturbations (Theodoris et al., 2023). State-feedback ISP passes the model’s inferred response from one step to the next, so that the effect of a perturbation can depend on the cell’s response to earlier ones, and its placebo-contrasted effect size separates factor-specific feedback from the gene-independent drift that feedback introduces (Fig. 4A, B).

Under feedback, the order of the Yamanaka factors changed their factor-specific effect: adding POU5F1 or SOX2 first and KLF4 or MYC last gave the largest effects, and only these orders exceeded simultaneous OSKM (Fig. 4C, D). This is an effect within the model, in which a schedule is an order of perturbations of one cell’s gene ranking rather than a time course of factor delivery. It should not be read as a prediction of the optimal temporal sequence of factor delivery. In particular, the schedule [POU5F1, KLF4]→MYC→SOX2, which outperformed simultaneous delivery in mouse fibroblasts through a sequential EMT and MET mechanism (Liu et al., 2013), did not exceed simultaneous OSKM in our analysis. Whether feedback within the model captures the temporal requirements observed experimentally remains to be tested, for example against time-resolved reprogramming data.

State-feedback ISP has further limitations. The reordering applied at each feedback event grew over successive steps (1.9-fold between the first and the fourth event), so errors may accumulate in long schedules. The size of τ depended on the number of cells used to train the decoder, so τ values should be compared only across analyses that use the same decoder training settings. By contrast, how clearly OSKM separated from the placebos (for example, z) did not depend on those settings. Because each schedule was scored with its own decoder, differences in τ between schedules mix the effect of factor order with differences between decoders.

Targeted NANOG-plus-partner screens extended the cross-species comparison to the choice of goal state in primed human iPSCs. Partner prioritization was highly goal-dependent: DPPA3 ranked highest toward the human P20N naïve goal, whereas ESRRB ranked highest toward the mouse DiPSC_2i ground-state reference, and DPPA3 and TFCP2L1 ranked high toward both (Fig. 5A, C). Because these goal embeddings probe related but non-identical axes, their moderate rank concordance (Spearman ρ ≈ 0.35) indicates that the P20P→DiPSC_2i trajectory likely captures a complex mixture of pluripotency and species-specific signatures. Both rankings were nevertheless led by naïve-associated factors (DPPA3 and ESRRB), so ISP recovered the shared pluripotency program under either goal. The differences between the rankings reflect the other differences between the start and goal cells that each goal embedding also contains, so the goal state should be chosen to match the biological program of interest.

These observations highlight important considerations for formulating hypotheses and interpreting ISP results. A small predicted shift does not necessarily preclude wet-lab experimental efficacy. Rather, it may reflect a highly specific transition, such as an early epigenetic unlocking (Buganim et al., 2012; Polo et al., 2012), that aligns poorly with the global transcriptome of the defined goal state (e.g., the modest shift induced by the Yamanaka factors in Fig. 3B). Conversely, a large predicted shift does not guarantee the acquisition of an intended phenotype (such as the predicted shift from the primed toward the naïve state in Fig. 5B), but may instead capture an intermediate transition state or transient localized remodeling along the trajectory. Thus, the ISP Platform, which makes Cross-species, State-feedback, and Multi-gene ISP available through a graphical user interface, is best utilized not as an absolute binary predictor, but as a hypothesis-generating framework that requires careful formulation of goal states and context-aware biological interpretation to guide focused experimental follow-up.

## 4. Experimental Procedures

### 4.1 ISP Platform and workflow

All analyses were performed with ISP Platform (v1.0.2) (https://github.com/YuyaSanaki/ISP-Platform). Development and computational runs were executed on DGX Spark (NVIDIA) or Pegasus supercomputer (NVIDIA H100 GPUs) at the Center for Computational Sciences, University of Tsukuba.

### 4.2 Model interchangeability architecture

We decoupled the input species from the inference model, enabling human and mouse Geneformer variants to be used interchangeably (Chen et al., 2026; Ito et al., 2025; Theodoris et al., 2023). Each run specified the input species and Geneformer variant, which determined the checkpoint, token dictionary, and maximum input length (4,096 and 2,048 tokens for human and mouse models, respectively). Matched inputs were directly tokenized, whereas swapped-model inputs underwent orthology-aware identifier conversion, target-model normalization, and rank-value re-tokenization. Orthologs between human and mouse were retrieved from Ensembl BioMart (release 116, human GRCh38.p14 and mouse GRCm39), and these tables are distributed with the ISP Platform v1.0.2 together with their SHA-256 checksums. Users can replace them with the current Ensembl release through an installation option. In orthology-aware identifier conversion, gene symbols were resolved to source-species Ensembl IDs (version suffixes removed), and each gene was replaced by its target-species ortholog. By default, only pairs annotated as one-to-one by Ensembl and unique in both directions were retained. One-to-many, many-to-one and many-to-many relationships were discarded, so that expression was never summed across paralogues, which would distort the rank-value encoding. The curated table of the ISP Platform then restored established pairs that Ensembl labels one-to-many (POU5F1, NANOG and GAPDH, as of v1.0.2). The human paralogues POU5F1B and NANOGP8 remained unmapped. Genes without a retained ortholog were removed. In target-model normalization and rank-value re-tokenization, the ISP Platform recalculated the expression of the ortholog-converted genes in the target model’s terms. Following the original Geneformer implementation (Chen et al., 2026; Ito et al., 2025; Theodoris et al., 2023), counts of the genes in the target model’s vocabulary were normalized to each cell’s total over these genes and divided by the target model’s gene-specific medians, so that cross-species input was normalized in the same way as native input of the target model. The genes were then re-ranked by these values and encoded with the target model’s vocabulary and input size. To monitor ortholog conversion, the ISP Platform reports the mapping yield, the fraction of input genes converted to target-species orthologs. In this study, the yield was 48.9% for human-to-mouse conversion and 57.6% for mouse-to-human conversion. The removed genes were mostly non-coding or rarely detected. To avoid losing genes of interest, the optional ortholog-loss gate stops a run before tokenization when any user-specified critical gene is dropped. Users can resolve this stop by supplying a project-level curated table of ortholog pairs, specified in the Web UI or the configuration file (a template is provided in examples/ortholog_overlays/). Users can also request that established orthologs be added to the platform curated tables through GitHub issues.

### 4.3 Datasets, tokenization, and fine-tuning

We analyzed public single-nuclei and single-cell transcriptomic datasets of fibroblast-to-iPSC reprogramming: human GSE147564 (Liu et al., 2020) and mouse GSE115943 (Schiebinger et al., 2019). Somatic cells were defined as day-0 fibroblasts (D0 fibroblasts in human, D0_Dox MEFs in mouse). In the human dataset, P20P and P20N denote passage-20 iPSCs maintained under primed and naïve pluripotency conditions, respectively. In the mouse dataset, DiPSC_2i was used as the 2i-maintained ground-state iPSC reference. Raw count matrices were filtered with a quality-control policy (Supplementary Table 1), retaining 3,000 somatic cells and 3,000 iPSCs (P20P in human, DiPSC_2i in mouse) per analysis arm. The cells were tokenized with the dictionary of the matched or swapped model. Tokenization, fine-tuning, and ISP used the Geneformer code (Theodoris et al., 2023) as adapted for Mouse-Geneformer (Ito et al., 2025). The human model was Geneformer V2-104M (Chen et al., 2026), and the mouse model was the six-layer Mouse-Geneformer (Ito et al., 2025). Inputs to Geneformer V2 (Chen et al., 2026) were encoded without the `<CLS>` and `<EOS>` tokens used in its pretraining, and all layers were fine-tuned on these encodings.

In each of the four arms (human or mouse cells, human or mouse model), the pretrained model was fine-tuned to classify somatic cells and iPSCs. Of the cells, 80% were used for training and 20% for evaluation. Fine-tuning ran for one epoch with a learning rate of 5 × 10⁻⁵ (linear decay after a 5% warm-up) and a weight decay of 0.001. For the primed-to-naïve screens, 3,000 P20P and 3,000 P20N cells were selected with the same quality-control policy, and the human model was fine-tuned on them in the same way. For the route toward the mouse ground state, the human model was fine-tuned in the same way on P20P cells and DiPSC_2i cells converted to human orthologs. Each analysis used its corresponding fine-tuned checkpoint.

A cell embedding was the mean of the final-layer hidden states over the genes of the cell (mean pooling, padding excluded). For visualization, cell embeddings were reduced to 50 principal components and projected by UMAP (McInnes et al., 2018) (Fig. 2A, random seed 0).

### 4.4 *In silico* perturbations

*In silico* overexpression (OE) and knockdown (KD) were implemented by manipulating the rank-value encoding of genes represented in each model, as in Geneformer (Theodoris et al., 2023). For OE, the target-gene token was inserted at the front of the rank-value encoding, corresponding to the highest expression rank (Theodoris et al., 2023). Geneformer inserts a gene that is absent from the encoding and truncates the encoding only when it exceeds the model input size. In the ISP Platform, each inserted gene instead displaced the lowest-ranked gene, so that every cell kept its original length. For KD, the corresponding token was removed from the encoding (deletion in Geneformer, Theodoris et al., 2023). In multi-gene OE, genes already in the encoding were moved to the front, absent genes were inserted at the front, and one lowest-ranked gene was removed for each inserted gene. Before embeddings were compared, overexpressed genes and genes not shared between the original and perturbed encodings were excluded, and the remaining embeddings were matched by gene identity (Fig. 3A).

Perturbation effects were quantified as the Geneformer goal-state shift (Theodoris et al., 2023). For each cell, the shift is the cosine similarity between the perturbed cell embedding and the goal-state embedding (the mean embedding of the goal-state cells), minus that of the unperturbed cell embedding. A positive shift indicates movement toward the goal state. In genome-wide screens, the shift of each gene (“Shift_to_goal_end”) was the mean over the cells whose encoding contains the gene, and the number of these cells is given as N_Detections. Statistical significance in genome-wide screens was assessed using the false discovery rate of Geneformer (Theodoris et al., 2023), which compares the shifts of each gene with those of all perturbed genes (Wilcoxon rank-sum test, Benjamini and Hochberg correction). For multi-gene OE, the median of the per-cell shifts is reported, and conditions applied to the same cells were compared by two-sided paired Wilcoxon signed-rank tests with Bonferroni correction.

For ISP UMAP visualizations, the start cells, the same cells after perturbation, and the goal-state cells were embedded together and projected by UMAP in the same way. Lines connect the positions of 50 randomly chosen cells before and after perturbation. Their length is a distance in the two-dimensional UMAP plane and is not calibrated to the goal-state shift.

### 4.5 Genome-wide KD screens and pathway enrichment

Genome-wide KD screens started from the 3,000 iPSCs of each arm, with the somatic cells as the goal state, in the two matched and the two swapped-model arms (Fig. 2B). In every cell, each gene of the encoding was deleted in turn. Hits were filtered by FDR < 0.05, positive Shift_to_goal_end, and N_Detections > 20, and ranked by Shift_to_goal_end. Pathway enrichment of the top 100 hits of each arm was tested with Enrichr (Kuleshov et al., 2016; Xie et al., 2021) through GSEApy (Fang et al., 2023) against the Reactome Pathways 2024 library (Milacic et al., 2024) (Fig. 2C). The background was all genes tested in the screen, and gene symbols were those of the model species. Terms with FDR < 0.05 (Benjamini and Hochberg correction) were regarded as enriched.

### 4.6 Multi-gene OE of reprogramming factors and NANOG-plus-partner screens

Multi-gene OE of OSKM, OSN, and the 7-factor cocktail was applied to 1,000 somatic cells with the human-matched model, with P20P iPSCs as the goal state (Fig. 3B). For the ISP UMAP (Fig. 3C, D), a separate set of 3,000 somatic cells and 3,000 P20P iPSCs was used (random seed 7).

Targeted two-gene OE screens started from 3,000 P20P cells with the human model. The goal state was either P20N cells or DiPSC_2i cells. DiPSC_2i cells were converted to human orthologs and tokenized together with the P20P cells in the human vocabulary. Each route used the checkpoint fine-tuned on its start and goal cells. NANOG was overexpressed together with each of 21 candidate partners, with KLF2 as the benchmark partner (Takashima et al., 2014), and with each of four negative-control genes (DDO, KRTAP12-3, ASIC5, and PDHA2). The candidate partners were known pluripotency factors: naïve-associated transcription factors (ESRRB, GBX2, KLF4, KLF5, KLF17, PRDM14, TBX3, and TFCP2L1), naïve pluripotency markers (DPPA2, DPPA3, DPPA4, and DPPA5), core pluripotency factors (FOXD3, LIN28A, MYC, POU5F1, SALL4, SOX2, UTF1, and ZFP42), and STAT3. NANOG and KLF2 were also overexpressed alone. The baseline aggregate was the mean of the median shifts of the four negative-control pairs. The partner rankings of the two routes were compared by Spearman correlation among the partners tested on both routes. ISP UMAPs of the top partner of each route (Fig. 5B, D) used 3,000 cells per group (random seed 0).

### 4.7 State-feedback ISP

State-feedback ISP applies perturbations in ordered steps and feeds the model’s response back after every step (Fig. 1C). Each step applies the same rank edits as single-step ISP to the current encoding, and a gene overexpressed in a later step is placed in front of those overexpressed earlier. Geneformer has no output for expression. A linear Δrank decoder therefore translates the model’s response into a new gene order. Its input is, for each gene, the change in final-layer hidden states caused by the perturbations so far (perturbed minus original start encoding, matched by gene). Its output is a displacement of the gene’s rank. The decoder is a single linear layer without bias, bounded by a hyperbolic tangent and a maximum displacement, so that no change in hidden state gives no displacement. After each step, the genes of the cell were reordered by their original rank plus the predicted displacement, and the reordered encoding was the input to the next step. Genes overexpressed so far were kept at the front, and no gene was added or removed by the reordering.

A decoder was trained for each schedule and decoder seed. The decoder was trained on the start cells of the analysis (300 cells in Fig. 4A, B and 100 cells in Fig. 4C–F), with up to 256 genes per cell other than the factors. The teacher was the observed rank displacement of each gene between pseudobulk somatic and pluripotent cells, that is, the difference in the gene’s mean position divided by the mean encoding length of the state, over up to 3,000 cells of each state. Genes detected in fewer than five cells of either state were not used. The training input was the hidden-state change with all steps of the schedule applied at once. Genes, not cells, were split 80:20. The decoder was trained on 80% of genes with Huber, pairwise ordering, and smoothness losses. The maximum displacement (0.05, 0.1, 0.2, 0.3, or 0.5 of the encoding length) was chosen by the Spearman correlation between predicted and observed displacement on the held-out 20%. The goal-state shift at the end point was not used to train or select the decoder.

The goal-state shift after each step was computed against the original start encoding. Because the reordering does not keep genes at fixed positions, cell embeddings were mean-pooled over all genes of each encoding, including the overexpressed genes, without removing positions from the unperturbed encoding.

Part of the shift produced by feedback also appears when other genes are added in place of the factors, so it is not specific to the factors. This non-specific shift was removed with two groups of placebo gene sets, correction placebos and comparison placebos, drawn in the same way with different random seeds. Correction placebos were used within the feedback loop in a potential-outcome contrast, and comparison placebos were run through the whole procedure and their effect was subtracted from that of the factors (Δspec). Placebo gene sets were drawn from the whole vocabulary of the model, excluding special tokens and the factors. Each placebo gene was matched to one factor in the start cells. For a factor absent from all start cells, placebo genes were drawn from genes absent from all start cells. Otherwise they were drawn from genes detected in half to twice as many start cells and with a median normalized position within 0.1 of the factor’s. This window was widened to 0.2, 0.3, or no limit when fewer than 50 genes qualified. A placebo chain replaces each factor by its matched placebo gene in the same step. The placebo gene sets are listed in Supplementary Table 2.

In the potential-outcome contrast, the 10 correction placebo gene sets were swapped in for the factors in the current encoding at each step. Their mean hidden-state change estimates how each gene would change under the same perturbation with genes other than the factors, and it was subtracted from the hidden-state change caused by the factors before decoding. The decoder was trained on the same contrast.

The feedback gain of a cell was its end-point shift minus the end-point shift of the same steps applied without feedback. The comparison placebo chains were run through the whole procedure under the same conditions, with the decoder trained for the schedule. The factor-specific effect τ (Δspec) of a cell was the feedback gain of the factors minus the mean feedback gain of the comparison placebo chains in the same cell. τ of a schedule is the mean over cells, with a 95% confidence interval from 2,000 bootstrap resamples of cells. Separation from the placebos was summarized as z = (τ_OSKM − mean τ_placebo) / SD τ_placebo, with τ of each placebo chain computed in the same way. Analyses were repeated with decoder seeds 0, 1, and 2.

To choose the correction, OSKM was added in the order KLF4→MYC→SOX2→POU5F1 to 300 somatic cells, with P20P iPSCs as the goal state, and compared with 30 comparison placebo sets (Fig. 4A, B). Six methods were applied to the same cells: no correction (decoder trained and applied on the hidden-state change itself), Δspec only, backdoor adjustment (removes baseline bias: hidden-state change minus its ridge-regression prediction from the hidden state in the start encoding, fitted on the correction placebos, with a decoder trained on the adjusted change), mediator adjustment (removes the non-specific response to the rank edit: the uncorrected decoder applied to the hidden-state change minus the mean change of the correction placebos swapped in for the factors in the current encoding), potential-outcome contrast only, and potential-outcome contrast plus Δspec. Backdoor adjustment, mediator adjustment, and the potential-outcome contrast were applied at every feedback step. The share of the placebo response was the mean feedback gain of the placebo sets relative to that of OSKM.

For the comparison of schedules, the 26 schedules (the 24 orders that add one factor per step, [POU5F1, KLF4]→MYC→SOX2, and simultaneous OSKM) were run on the same 100 somatic cells with decoder seeds 0–2 (Fig. 4C, D). Placebo chains replaced each factor in its step by a gene with the same detection and rank stratum in the start cells, and the same 20 draws were used for every schedule. The agreement of the schedule rankings between decoder seeds was measured by Spearman correlation. Schedules were compared with simultaneous OSKM by two-sided paired Wilcoxon signed-rank tests on per-cell τ averaged over seeds, with Holm correction over 25 schedules.

For the highest-ranked order, the goal-state shift after the feedback of each step was compared with that of its 20 placebo chains (Fig. 4E). To show these steps in embedding space (Fig. 4F), principal component analysis was fitted on the embeddings of 1,000 reference somatic cells and 1,000 reference P20P iPSCs, and the cells before perturbation and after the feedback of each step (decoder seed 0) were projected onto the first two components. The position of each cell between the somatic centroid (0) and the iPSC centroid (1) was computed in the full embedding space by projecting the cell onto the line through the two centroids.

## Supporting information

supt1

supt2

## Acknowledgments

We are grateful to Hiromi Yanagisawa and Marina Sanaki-Matsumiya for valuable technical suggestions. Authors used Gemini, Perplexity, and Cursor to assist with text editing and coding.

## Declarations

### Author Contributions

**Yuya Sanaki**: Conceptualization, Project administration, Formal analysis, Funding acquisition, Investigation, Validation, Visualization, Writing – original draft. **Shuji Shigenobu**: Methodology, Resources, Writing – review & editing. **Ryusuke Niwa**: Funding acquisition, Project administration, Writing – review & editing.

### Data Availability

Raw data are available from GEO (GSE147564, GSE115943). Count matrices used in this study (QC and subsampled) are deposited in the University of Tsukuba Repository (https://doi.org/10.15068/0002025690).

The ISP Platform and related materials used in this study, including the Ensembl ortholog tables and analysis code, are available at https://github.com/YuyaSanaki/ISP-Platform (version 1.0.2).

