## Supplementary material for "ISP Platform powered by Geneformer: Framework for Cross-Species, State-Feedback, and Multi-Gene *In Silico* Perturbation Screens with Application to iPS Cell State Transitions": supt2

### Supplementary Table 2. Placebo gene sets of State-feedback ISP (Fig. 4)

Each placebo gene replaces one Yamanaka factor and is overexpressed in the step of that factor. Placebo genes were drawn at random from the vocabulary of the human Geneformer model, excluding special tokens and the four factors, and without replacement across sets. Each gene was matched to its factor in the start cells. It was detected in half to twice as many start cells as the factor, and its median normalized position was within 0.1 of the factor's (widened to 0.2, 0.3 or no limit when fewer than 50 genes qualified). POU5F1 and SOX2 were absent from all start cells, so their placebo genes were drawn from the genes absent from all start cells.

Correction sets (C1 to C10) were subtracted within each step in the potential-outcome contrast. Comparison sets (P) were run through the whole procedure and give τ of the placebos. The two groups were drawn with different random seeds, so a gene can occur in both. A correction set that shares a gene with a comparison set is left out of the contrast of that set.

(A) Fig. 4A, B: 300 somatic cells, the same sets for decoder seeds 0, 1 and 2. (B) Fig. 4C–F: 100 somatic cells, the same sets for all 26 schedules and decoder seeds. In a placebo chain, each factor is replaced by its placebo gene in the same step. (C) Matching of each factor. Detection, fraction of start cells whose encoding contains the factor. Median position, median of the factor's position divided by the encoding length minus one, over the start cells that contain it (0, first rank). Window, largest allowed difference in median position. Genes, number of genes in the stratum from which its placebo genes were drawn.

#### (A) Fig. 4A, B (300 cells)

| Group | Set | POU5F1 | SOX2 | KLF4 | MYC |
|---|---|---|---|---|---|
| Correction | C1 | SDR9C7 | SPINK9 | RIPOR2 | AKR1C3 |
| Correction | C2 | CRYBA4 | SSX2B | LRP1B | ANGPTL4 |
| Correction | C3 | POM121L2 | TREML4 | TNFRSF1B | ADAM23 |
| Correction | C4 | TMEM238L | C4orf50 | PDE1A | EPHA4 |
| Correction | C5 | ENSG00000267173 | UGT1A1 | SAMHD1 | VPS13A |
| Correction | C6 | LYPD6B | RAB9B | RPL38 | EYA4 |
| Correction | C7 | ENSG00000291954 | SALL2 | SNCA | DDIT4 |
| Correction | C8 | NAT8 | BHMT | S100A2 | TRPM3 |
| Correction | C9 | ENSG00000282034 | MFRP | SLC1A3 | TNRC6C |
| Correction | C10 | SPON1 | CARD11 | ITPKB | SCD5 |
| Comparison | P1 | TP53I13 | OR4D9 | RPS21 | NLRC5 |
| Comparison | P2 | GAGE12J | CCK | SRRM3 | CACNA1A |
| Comparison | P3 | PTGR2 | SCN4B | KRT81 | COX6C |
| Comparison | P4 | PSORS1C2 | TAAR9 | CDA | NR4A1 |
| Comparison | P5 | SPEM3 | F8A3 | GPC5 | IFI6 |
| Comparison | P6 | PDX1 | ENSG00000290149 | ATP10B | ADGRL4 |
| Comparison | P7 | VSX2 | ENSG00000111780 | MFAP5 | DOCK9 |
| Comparison | P8 | SPDYE14 | GPR25 | KIAA1755 | PATJ |
| Comparison | P9 | ENSG00000288236 | SPRR1B | BTN3A2 | DPT |
| Comparison | P10 | CLDN7 | DLGAP2 | KIF21A | ARF6 |
| Comparison | P11 | C12orf54 | TREM2 | DLGAP1 | SRSF10 |
| Comparison | P12 | CASR | CD300E | RPS12 | GJA1 |
| Comparison | P13 | PDCD6-AHRR | ENSG00000284626 | CPNE4 | NDST3 |
| Comparison | P14 | RNASE7 | ENSG00000285558 | IDS | GBP1 |
| Comparison | P15 | DPPA4 | EXOSC3 | CSMD1 | RGS19 |
| Comparison | P16 | HSPE1 | ENSG00000284732 | NCAM2 | LMOD1 |
| Comparison | P17 | ENSG00000259171 | ENSG00000277629 | LRRC7 | ANGPTL4 |
| Comparison | P18 | ARMH2 | CIROP | TIAM1 | CCND3 |
| Comparison | P19 | SSMEM1 | OR2L5 | CADM2 | IRF1 |
| Comparison | P20 | BCHE | KIR3DL2 | EFEMP1 | SOCS3 |
| Comparison | P21 | TNFSF13 | SSX5 | CTSH | CAMK4 |
| Comparison | P22 | ENSG00000275822 | OR2J1 | TBX20 | SERPING1 |
| Comparison | P23 | ENSG00000286239 | OR5AK3P | ACAP1 | PDE3A |
| Comparison | P24 | CTAGE15 | H2BC26 | LMOD1 | IFI44 |
| Comparison | P25 | ATXN3L | CHL1 | TGM2 | FRY |
| Comparison | P26 | FAM90A9 | SMKR1 | RPS27 | IMPDH2 |

| Group | Set | POU5F1 | SOX2 | KLF4 | MYC |
|---|---|---|---|---|---|
| Comparison | P27 | RAB3A | GOLGA7B | THSD7A | ARHGAP26 |
| Comparison | P28 | ENSG00000273154 | PRODH | LRMDA | EGR1 |
| Comparison | P29 | EID3 | ENSG00000275778 | MSR1 | SLCO3A1 |
| Comparison | P30 | ENSG00000276256 | RNF122 | KCNIP4 | ROMO1 |

#### (B) Fig. 4C–F (100 cells)

| Group | Set | POU5F1 | SOX2 | KLF4 | MYC |
|---|---|---|---|---|---|
| Correction | C1 | GPT | GPX6 | FGF23 | SRGAP3 |
| Correction | C2 | SPATA31F3 | GFY | H1-4 | ARHGAP42 |
| Correction | C3 | ARL11 | NDUFA13 | TNFRSF1B | MARK1 |
| Correction | C4 | ENSG00000258466 | SUMO4 | F13A1 | MFAP2 |
| Correction | C5 | ARMH2 | PRAMEF6 | CTSH | TAFA2 |
| Correction | C6 | TIMD4 | TEX11 | MCTP1 | RNF130 |
| Correction | C7 | ENSG00000291851 | PROK2 | ITPR1 | ANGPTL4 |
| Correction | C8 | RLBP1 | VAV1 | MAFB | PSMC5 |
| Correction | C9 | C17orf78 | TMEM233 | RBFOX1 | OSBPL6 |
| Correction | C10 | SPMIP11 | HLA-DRB1 | OCIAD2 | SFRP2 |
| Comparison | P1 | FBP2 | PROB1 | SERPINB2 | NR4A2 |
| Comparison | P2 | FXYD3 | LAD1 | GAD1 | C4orf48 |
| Comparison | P3 | GFAP | ZNF793 | NECAB1 | IL2RA |
| Comparison | P4 | ENSG00000274053 | FAM9B | MT1A | NRCAM |
| Comparison | P5 | MS4A5 | SLC34A3 | SORBS2 | APOBEC3G |
| Comparison | P6 | CPA1 | CHGB | HCN1 | SAMD3 |
| Comparison | P7 | ALPK3 | LHX8 | GBP4 | LYN |
| Comparison | P8 | SRARP | MYLK3 | AQP1 | RUNX3 |
| Comparison | P9 | HLA-DRB4 | CNKSR1 | ZMAT4 | NDUFAB1 |
| Comparison | P10 | SMTNL2 | AKR7L | PSCA | GAS7 |
| Comparison | P11 | SPANXC | ITGAD | GALNT18 | ARRDC3 |
| Comparison | P12 | CCL15-CCL14 | GARIN6 | OPCML | UNC5D |
| Comparison | P13 | HOMEZ | DAZ2 | GABRB3 | GALNT13 |
| Comparison | P14 | ENSG00000284111 | STOX1 | LTF | PATJ |
| Comparison | P15 | ENSG00000275117 | CLDN5 | SLC5A11 | SAMD5 |
| Comparison | P16 | DMRTC1B | USP43 | ARHGAP15 | EPB41 |
| Comparison | P17 | WDR83OS | GOLGA6L1 | ANKS1B | CDK1 |
| Comparison | P18 | STMND1 | CGB2 | CNTNAP5 | DNAJB1 |
| Comparison | P19 | DRD2 | AGT | MCTP1 | RESF1 |
| Comparison | P20 | PERCC1 | AGBL4 | NEDD4L | COX17 |

#### (C) Matching of each factor

| Analysis | Factor | Detection | Median position | Window | Genes |
|---|---|---|---|---|---|
| Fig. 4A, B (300 cells) | POU5F1 | 0.000 | absent | not applicable | 7,116 |
| Fig. 4A, B (300 cells) | SOX2 | 0.000 | absent | not applicable | 7,116 |
| Fig. 4A, B (300 cells) | KLF4 | 0.037 | 0.918 | 0.1 | 259 |
| Fig. 4A, B (300 cells) | MYC | 0.073 | 0.864 | 0.1 | 333 |
| Fig. 4C–F (100 cells) | POU5F1 | 0.000 | absent | not applicable | 8,133 |
| Fig. 4C–F (100 cells) | SOX2 | 0.000 | absent | not applicable | 8,133 |
| Fig. 4C–F (100 cells) | KLF4 | 0.020 | 0.935 | 0.1 | 371 |
| Fig. 4C–F (100 cells) | MYC | 0.060 | 0.876 | 0.1 | 384 |
